# Probing Causality of the Brainstem-Hypothalamic Murine Models of Sleep-Wake Regulation

**DOI:** 10.1101/2020.09.21.306456

**Authors:** Fatemeh Bahari, Myles W. Billard, John Kimbugwe, Carlos Curay, Glenn D.R. Watson, Kevin D. Alloway, Bruce J. Gluckman

## Abstract

Sleep-wake regulation is thought to be governed by interactions among several nuclei in midbrain, pons, and hypothalamic regions. Determination of the causal role of these nuclei in state transitions requires simultaneous measurements from the nuclei with sufficient spatial and temporal resolution. We obtained long-term experimental single- and multi-unit measurements simultaneously from multiple nuclei of the putative hypothalamic and brainstem sleep-wake regulatory network in freely behaving rats. Cortical and hippocampal activity, along with head acceleration were also acquired to assess behavioral state. Here, we confirm that the general activity profile of the recorded sleep-wake regulatory nuclei is similar to the patterns presented previously in brief recordings of individual nuclei in head-fixed animals. However, we found that these activity profiles when studied with respect to cortical and behavioral signs of state transitions differ significantly from previous reports. Our findings pose fundamental questions about the neural mechanisms that maintain specific states and the neural interactions that lead to the emergence of sleep-wake states.

## 1. Introduction

Sleep-wake states are thought to be driven by basal forebrain, brainstem, and hypothalamic circuits (ECONOMO, 1930; Moruzzi and Magoun, 1949; Saper et al., 2005, 2010; Datta and MacLean, 2007; Weber et al., 2015; Xu et al., 2015; Sherin et al., 2018). These states are predominantly characterized from electroencephalogram (EEG) and electromyogram (EMG) recordings, with transitions between states described as discrete mechanisms (Saper et al., 2001, 2010; Datta and Siwek, 2002; Luppi et al., 2006; Weber and Dan, 2016). Models of underlying network mechanisms of sleep-wake regulation have attempted to describe and reproduce these discrete transitions (Tamakawa et al., 2006; Fulcher et al., 2008; Phillips and Robinson, 2008; Diniz Behn and Booth, 2010; Rempe et al., 2010; Fleshner et al., 2011). To do so, these models often invoke co-inhibitory dynamics of sleep-wake regulatory cell groups in configurations analogous to electrical flip-flops (Saper et al., 2001; Lu et al., 2006; Weber and Dan, 2016). However, observations of cell group activity and involvement are derived from experiments conducted on head-fixed (Boissard et al., 2002; Verret et al., 2003; Takahashi et al., 2010; Sakai, 2011; Boucetta et al., 2014) or lightly anesthetized animals (Aston-Jones et al., 2001; Sherin et al., 2018), via short recordings, and often within novel environments (Datta and Siwek, 2002; Lu, 2006; Alam et al., 2014; Weber et al., 2015; Xu et al., 2015; Bjorness et al., 2016). These conditions inherently modulate sleep-wake patterns (Sigl-Glöckner and Seibt, 2019). Further, short recording sessions provide too few state transitions to accurately determine the relationship between firing rate and state transitions. These recording sessions are often 2-6 hours long amounting to approximately 6-20 sleep cycles (Takahashi et al., 2009, 2010; Sakai, 2018). Following the Poisson distribution laws and a typical bin size of 10 ms for firing rate calculations, a few hundred state transitions are required to detect a statistically significant change in firing rates relevant to the within-state activities of these neurons.

To investigate the neural basis of sleep-wake transitions in normal conditions, we chronically recorded single- and multi-unit activity simultaneously from multiple hypothalamic and brainstem sleep-wake regulatory nuclei in freely behaving rats. Cortical and hippocampal activity, along with head acceleration were also acquired to assess behavioral state. The dataset comprises 100-300 sleep-wake cycles for each neuron, which provides a sufficient number of state transitions to detect a statistically significant change in firing rate. Our findings address current hypotheses regarding the causal role of the represented brainstem and hypothalamic cell groups in initiating naturally emerging transitions between states of vigilance.

## 2. Methods

All animal work was approved by and performed under the oversight of the Institutional Animal Care and Use Committee at the Pennsylvania State University.

To study the sleep-wake regulatory network as well as to identify sleep-wake transitions we used the systemDrive (Billard et al., 2018): a customizable micro-wire multitrode microdrive for targeting multiple cell groups along non-parallel non-co-localized trajectories. Animals were implanted with the systemDrive’s micro-wire multitrodes targeting several sleep-wake regulatory cell-groups. The systemDrive houses additional screw electrodes for electrocorticography (ECoG), and fixed depth electrodes for hippocampal local field potential (LFP) recordings. Following implantation, animals were returned to their home-cages and long-term continuous recordings started after ample post-operation recovery time (7 days).

In accordance with common microdrive system use, we advanced electrodes along the dorsal-ventral axis of each targeted cell group and recorded neuronal activity of the area for continuous periods of 3 to 10 days. The procedure was repeated until the entire depth of the target was covered. Single- and/or multi-unit activity, when present, was extracted for all channels in each target.

### 2.1. Animal Surgery and Care

Surgical and implantation techniques are previously described (Billard et al., 2018). Briefly, male and female Long Evans rats weighing between 275-350 grams were implanted for continuous recordings that lasted for a duration of 2-6 weeks.

Recording electrodes include micro-wire bundles for monitoring brainstem neuronal activity, as well as hippocampal LFP pairs and ECoG screws for determining the EEG and behavioral state. For observation of sleep-wake regulatory network (SWRN) dynamics, we recorded from five brainstem structures with target coordinates referenced to intraural line (IA) according to the Paxinos-Watson rat brain atlas:

Dorsal Raphe (DR) [AP: +1.5 mm, ML: 0 mm, DV: -3.6 mm, 21.33° or -21.33°]

PeduncloPontine Tegmentum (PPT) [AP: +0.5 mm, ML: ±2 mm, DV: -3.6 mm]

LateroDorsal Tegmentum (LDT) [AP: +0.36 mm, ML: ±0.8 mm, DV: -3 mm, 13.65° or -13.65°]

VentroLateral Preoptic Area (VLPO) [AP: +8.76 mm, ML: ±1 mm, DV: -0.75 mm, 15.52°]

Locus Coerleus (LC) [AP: -0.72 mm, ML: +1.3 mm, DV: -3.2 mm]

A number of these nuclei are located near sensitive structures such as the sagittal sinus and lateral ventricles. Therefore, trajectories were chosen to avoid these sensitive structures and allow enough room on the cranium for all electrodes.

For hippocampal LFPs, two pairs of custom-made 50 μm iridium oxide deposited micro-reaction chamber (μRC) electrodes formed from gold coated stainless-steel wire (Shanmugasundaram and Gluckman, 2017) with ends staggered by 250-300 μm were implanted bilaterally in the dorsal hippocampus at coordinates [AP: +5 mm, ML: ±3 mm, DV: -2.5 mm from the cortex]. For ECoG, four stainless steel screws (one-eighth inch 18-8 stainless steel 0-80 cortical screws, McMaster-Carr) were implanted at coordinates [AP: +7.5 mm, ML: ±4 mm], [AP: +3 mm, ML: -4 mm], and [AP: +2.5 mm, ML: +3 mm]. Two additional stainless screws were implanted at [AP: +11 mm, ML: ±3 mm] for ground and reference.

After completing surgery, animals were returned to their individual home-cages for recovery with free access to food and water. Following recovery, the animals were cabled for continuous video and EEG monitoring. Every 3 to 10 days the animals were briefly anesthetized to advance the micro-wire electrodes incrementally between recording sessions and until they covered the entire dorsal-ventral length of the structures of interest.

### 2.2. Histological Techniques

At the completion of recordings animals were sacrificed and their brains were histologically processed to examine the electrode tracks with respect to the targeted brain structures. Details and results of the histological methods and analysis are previously described (Billard et al., 2018).

### 2.3. Data Acquisition

All rats were housed individually in custom-made plexiglass cages with dimensions 9” (w) × 15” (d) × 24” (h). At the first electrode driving session, each animal received a differential or referential digitizing head-stage amplifier with 3-axis accelerometer (RHD2116 or RHD2132, Intan Technologies, Los Angeles, CA) which can directly connect to an electrode interface board on the systemDrive. The amplifier was connected to a data acquisition board (RHD2000 Evaluation System, Intan Technologies) via 3’ lightweight serial-peripheral interface (SPI) cable (Intan Technologies) and in-house adapted low-torque commutator (SRA-73540-12, Moog Inc.).

Hippocampal LFP, ECoG, as well as single and multi-unit signals were simultaneously acquired at 30,000 samples per second (SPS) and high-pass filtered at 1 Hz. Head acceleration measurements were sampled at a rate of 7500 Hz. A 12 hour light/dark cycle was maintained with the lights on between 6 am and 6 pm. Video monitoring, at 3 frames per second, began on the day of implantation and continued within each recording session. Video files were stored in one-hour-long files, while biopotentials and head acceleration data were stored in 5 to 15- minute-long files for further analysis.

To measure single- and multi-unit activity we used the microdrive arrangements within the systemDrive. For each animal, electrodes aimed at sleep-wake regulatory cell-groups in the brainstem and/or hypothalamus were first advanced after the recovery period.

During this and subsequent driving sessions, the animal was maintained under anesthesia with 0.5%-2% isoflurane gas and the head-mount was opened to access the systemDrive. We maintained the recordings throughout the driving session. Each electrode bundle was advanced incrementally (about 25 μm) and the signal was monitored until we observed new single-unit activity. For electrode trajectories that passed through the superior and inferior colliculi or other sensory-responsive regions, we tested auditory and visual stimuli to evoke responses that would confirm that the electrode was advancing through the appropriate brain structures.

### 2.4. Data Analysis

All analyses were performed using custom-written MATLAB (Mathworks) and Labview (National Instruments) scripts. Data used for analysis in this manuscript are available in an online repository.

#### 2.4.1. Sleep and Behavioral Scoring

Hippocampal LFP, ECoG, and accelerometer time-series were down-sampled to 1000 samples per second and reformatted into 1 hour-long blocks of binary data within a custom Labview script. The raw data were then band-pass filtered at 1-125 Hz for LFP, 1-55 Hz for ECoG, and 2-100 Hz for head acceleration.

The SOV was marked using an adaptation of the semi-automatic Linear Discriminant Analysis (LDA) previously described (Sunderam et al., 2007; Sedigh-Sarvestani et al., 2014). Briefly, the SOV was classified into five states: Non-rapid eye movement (NREM) sleep, rapid eye movement (REM) sleep, an intermediate state between NREM and REM (IS), awake (wake_wb_), and active exploration with non-coherent theta rhythm (wake_θ_). The frequency band specifications for NREM, REM, and awake states are as described in (Sunderam et al., 2007; Sedigh-Sarvestani et al., 2014). The IS was characterized by appearance of weak bursts of theta-band activity in the presence of identifiable delta band activity (GOTTESMANN, 1996; Boucetta et al., 2014). IS termination is defined by a large reduction of power in the lower delta band and a sharp increase in the theta band power (Fig. S1A). During the IS, the head acceleration power is low with no evidence of the short spikes characteristic of muscle twitches during REM.

For each animal, 4-6 hours of video-EEG data within 1 day were randomly selected and manually scored for SOV and SOV transition time. These data were then used to train the LDA. The remaining 18-20 hours of data were set aside as out-of-sample data (test set). For both training and test sets, features were computed, within 2 second long overlapping windows, from EEG spectral power in frequency bands 0.5-4, 4-8, 8-12, 12-25, and 25-80 Hz, plus coherence measurements (to distinguish REM bouts from IS and wake_θ_) and head acceleration. Head acceleration measurements were used for more accurate SOV scoring (Sunderam et al., 2007). The features were generated using causal filters and updated every second at the end of computation window. Therefore, all the scores are based on information up to the time leading to the state transition and *not* after that.

#### 2.4.2. Spike Scoring

We used Intan Technologies differential (RHD2216) and referential (RHD2132) preamplifiers to acquire electrophysiological signals. The default pass-band for the filters in these preamplifiers is between 0.1 Hz and 7.5 kHz. A 16-bit analog to digital converter then samples the signal from the AC-coupled amplifiers. At 7.5 kHz, the cut-off frequency in the analog low-pass filter is much higher than what is used conventionally (e.g. 3.5 kHz in (Datta and Siwek, 2002; Sakai, 2011)).

Data segments were filtered with a causal band-pass filter with 250 Hz to 7.5 kHz pass-band, and then thresholded to detect single-unit activity. Units were detected as instances of threshold crossings where thresholds were specified as multiples of the standard deviation of the filtered background signals. The background signals represent separate epochs of time that are free of distinguishable spiking activity and are used to calculate the mean and variance of the background distribution. We detected units with a threshold of 5-7 standard deviations. When spike events were detected, the waveform of each action potential was extracted from the data.

#### 2.4.3. Statistical Analysis

The core analysis of this work is to characterize the state-dependent firing rates and the peri- state-transition time firing rates. These measurements are calculated from the firing times of individual neurons and averaged over all like states or state-transition types. With just a mean-firing rate description, these measurement statistics should follow Poisson statistics, such that given a firing rate *FR*, the probability *P_λ_*(*k*) of observing the number of *k*; spikes measured within a time bin .1 with respect to the transition time summed over *N* observed transitions, as:

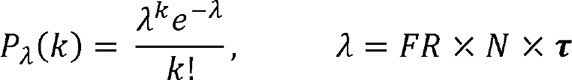

Critical in our analysis is to determine for each neuron when the instantaneous firing rate 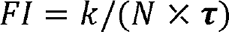 is significantly different from the pre-transition state mean firing rate, (see for example Fig. S3). We define the time *t_sig_* when the observed *k*; first falls within the upper or lower 1% of *P_λ_*(*k*).

During the course of our experiments, the activity of each neuron at each target location was measured over hundreds of sleep-wake cycles *N* є (100,300). Coupled with within-state and instantaneous firing rates typically much less than 20 per second, the large ensemble state transitions enables us to have sub-second resolution (*τ*) in our statistical analysis, which is crucial for causality determination.

## 3. Results

We successfully acquired simultaneous measurements from up to four separate sleep-wake regulatory nuclei (SWRN) in each rat. The success rates of chronic systemDrive targeting and recording from multiple (n = 12) rats are shown in Table S1. Cumulative recording duration, number of recording sessions, and number of single units exceeding a threshold of 7 standard deviations are presented for each animal. Data presented in this manuscript pertain to the SOV-dependent subset of neurons presented in Table S1.

Recording single units in deep brain structures across multiple days in freely moving rats is extremely challenging. The dataset described here was previously summarized in (Billard et al., 2018). In this previous work we confirmed the location of recorded targets using histological analysis (as shown in Fig. 6 in (Billard et al., 2018)). Further, to confirm long-term stability of our unit recordings we presented 1) measurements of signal-to-noise ratio for recordings from SWRN (Fig. 7 in (Billard et al., 2018)), 2) unit waveforms simultaneously extracted from up to 3 SWRN that were continuously recorded over 4-7 days from a subset of animals (Fig. 8 in (Billard et al., 2018)), and 3) examples of simultaneously recorded units from PPT, LDT, and DR over 6 continuous days (Fig. 9 in (Billard et al., 2018)). We therefore confirmed that unit detection is not disrupted during normal animal behavior and that recorded units remained stable throughout the duration of the recording periods with no significant deterioration of amplitude and signal-to-noise ratio.

**Figure 6.**
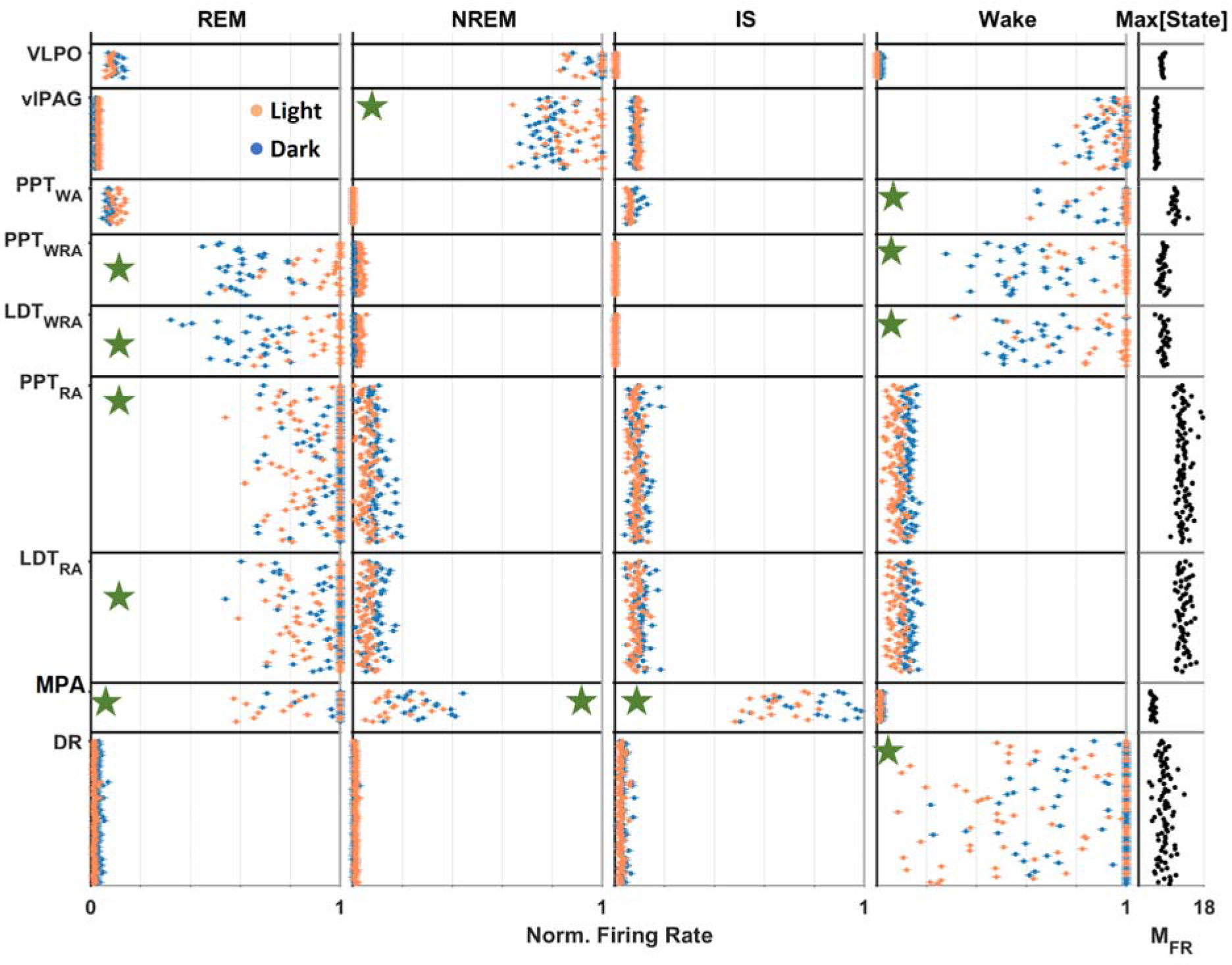
Mean and standard deviation of the firing rates of all recorded neurons in brainstem-hypothalamic sleep-wake regulatory network. Firing rates of state-dependent neurons whose recording sites were histologically confirmed pooled from all recording days across all animals. The neurons are separated based on their location and activity dependence. The firing rate of each neuron is marked by orange for the light period and blue for the dark period. For each neuron the firing rate is normalized to the maximum firing rate (black dots) of the group across all states of vigilance (M_FR_). Although the firing rates in all population were somewhat consistent, we found higher variability in subset of neuronal populations (green stars). In PPT and LDT we found a variety of neurons that were REM-active (RA), Wake and REM active (WRA), or Wake active (WA). VLPO, ventrolateral preoptical area; vlPAG, ventrolateral periacquaductal gray; PPT, pedunculopontine tegmentum; LDT, laterodorsal tegmentum; MPA, medial preoptic area; DR, dorsal raphe.

**Figure 7.**
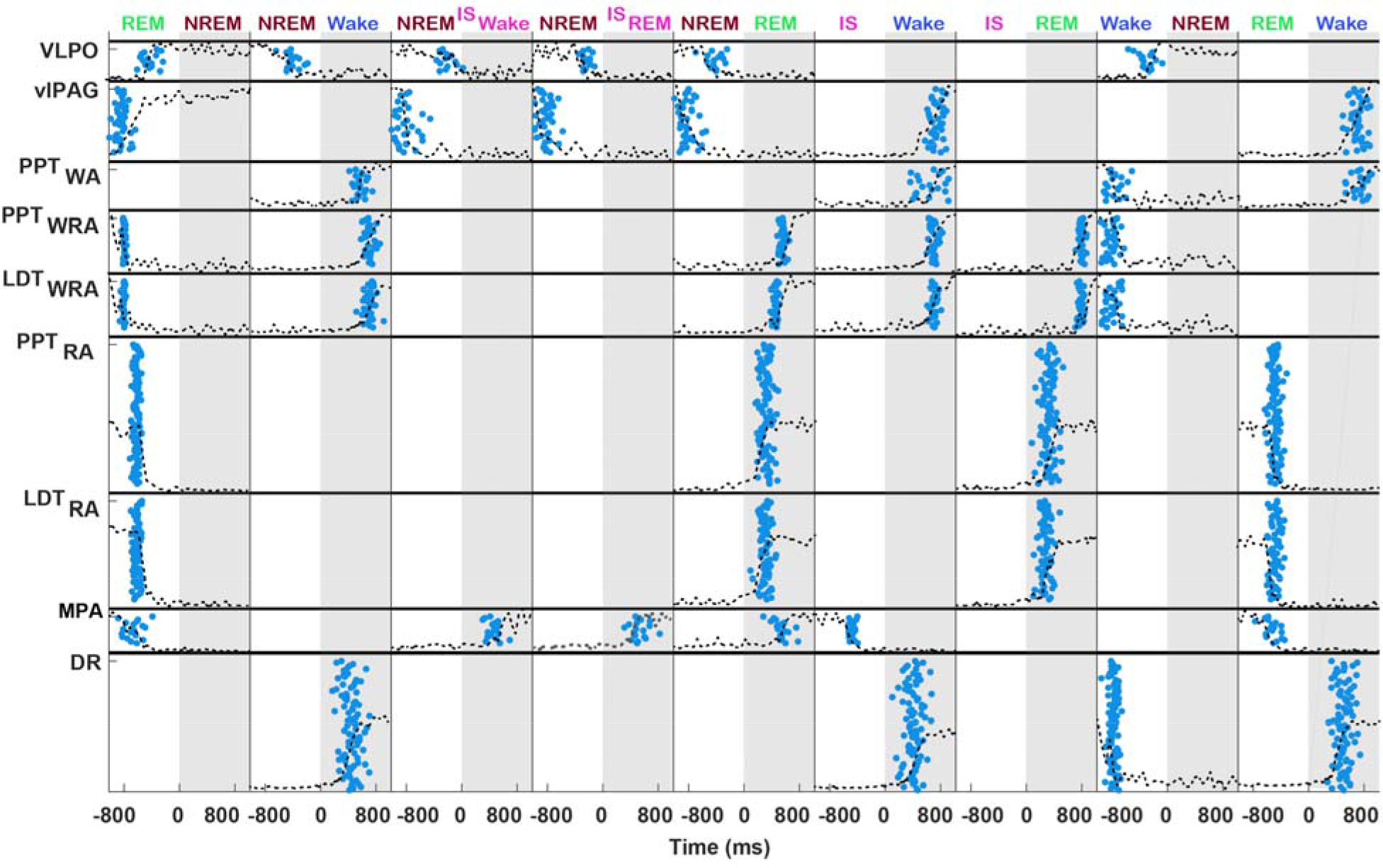
The t_sig_ for all neurons with respect to cortically defined state transitions. Each column indicates a state transition, where the transition is marked by time zero. The state before and after the transitions are marked by the white and gray backgrounds, respectively. The t_sig_ for each neuron is indicated by a blue dot. Representative average firing rates during transition are shown by the dashed gray lines. The firing rates for DR, PPT/LDT_RA_, PPT/LDT_WRA_ groups are normalized to half height of the plot box. Other firing rates are normalized to the full height of the plot box. Notice that for every neuronal group and transition, the t_sig_ values are tightly distributed. This indicates that the activity of these neurons is extremely consistent. The gray shading indicates the positive t_sig_ and not causal relationships while the white areas are negative t_sig_ indicating that the neuron changed its behavior prior to the appearance of cortical signs of a state. VLPO, ventrolateral preoptical area; vlPAG, ventrolateral periacquaductal gray; PPT, pedunculopontine tegmentum; LDT, laterodorsal tegmentum; MPA, medial preoptic area; DR, dorsal raphe.

We recorded single-unit activity with time-matched hippocampal, cortical, and head motion data scored for sleep-wake states. The dataset corresponds to 569 total days of recording spread over 12 animals. The SOV transition probabilities over all states are shown in Fig. S1D. The time-of-day dependence of the different SOVs are illustrated in Fig. S2A. This is likely the longest dataset acquired from freely behaving animals in their home cage environment for which the analysis includes identification of the intermediate state (IS). Although previous reports identify IS as the transitory state immediately prior to REM (GOTTESMANN, 1996; Gervasoni, 2004; Boucetta et al., 2014), we found that about 20% of the transitions out of IS continue into Wake (Fig. S1C&D). Additionally, the IS bouts in our recordings are typically 35-60 s long (Fig. S2B) which is much longer than previously reported.

### 3.1. Mean state-dependent neuronal firing rate and activity profiles of the brainstem-hypothalamic SWRN

In the brainstem-hypothalamic model of sleep-wake regulation interactions between cholinergic PPT/LDT, monoaminergic LC/DR, and GABAergic VLPO lead to different SOVs (Saper et al., 2010). Cumulatively, we recorded 855 single units across multiple regions and many continuous days (3-10 days) from 12 freely behaving animals all in their home-cage environment. A range of 100-300 complete sleep cycles were recorded from each neuron. About half (n = 428) of the identified single-units recorded were SOV-dependent Of the n=428 SOV-dependent neurons, 152 were REM-active, 15 were NREM active, 142 were wake-active, 60 were wake and REM active, and 41 were active in all SOVs except REM (denoted as REM-off). These neurons were distributed throughout multiple brain regions: We found Wake-active neurons in PPT, LDT, and DR; NREM-active neurons in VLPO; REM-active neurons in PPT and LDT. We also identified a group of REM-off neurons located in vlPAG. The anatomical distribution of the state-dependent and state-indifferent neurons is detailed in Table 1.

**Table 1.**
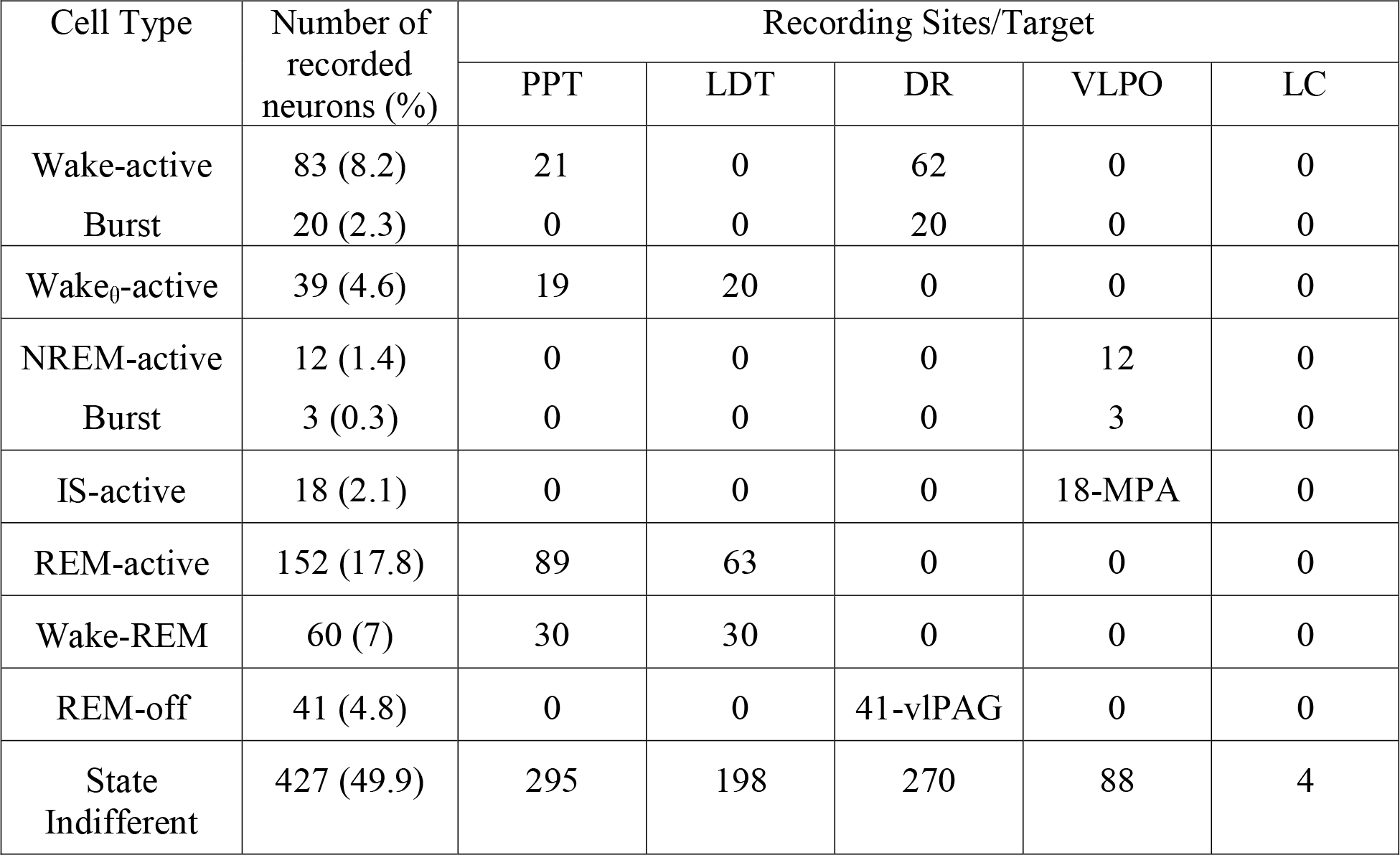
Anatomical distribution of the state-dependent and state-indifferent neurons in brainstem and hypothalamus. Listed are the number of neurons recorded in the indicated areas. In addition to the targeted brain regions, PPT, LDT, DR, VLPO, and LC, we measured state-dependent activity from adjacent structures. These include vlPAG from a subset of animals with electrodes targeting DR and neurons within the posterior region of LH. Abbreviations: PPT, pedunculopontine tegmentum, LDT, laterodosral tegmentum, DR, dorsal raphe, vlPAG, ventrolateral periacquaductal gray, VLPO, ventrolaterolateral preoptic area, MPA, medial preoptic area, LC, locus coerleus.

#### 3.1.1. Wake-active neurons

One hundred and forty two of the 428 state-dependent neurons (33%) were marked by having a higher discharge rate during wake periods than during sleep bouts. Of these, 20 – located in DR – displayed burst-like behavior with short inter-spike-interval (ISI) and relatively low firing rates (Fig.1). These neurons were specifically active during wake and not wake_θ_ bouts.

**Figure 1.**
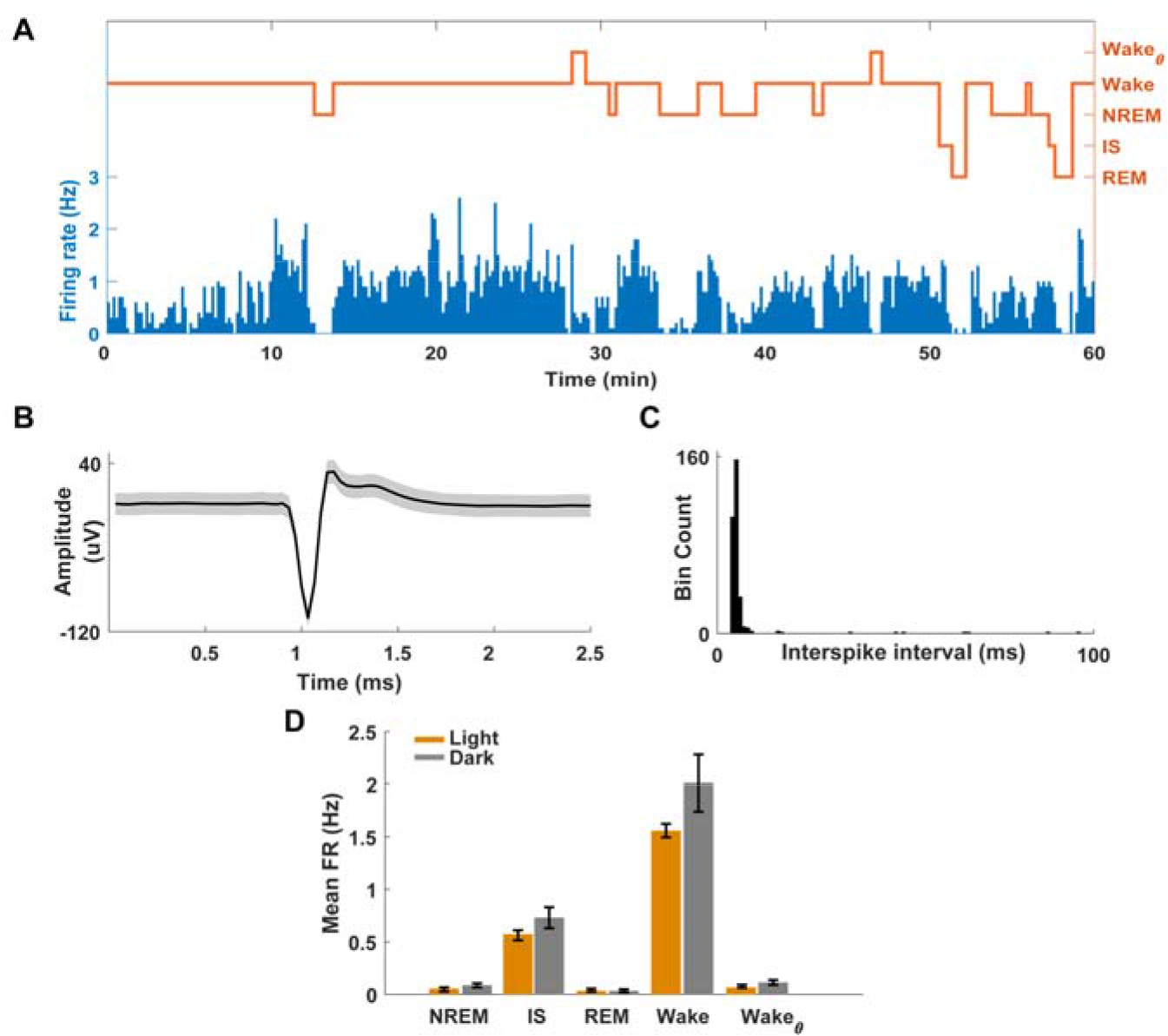
Wake-active neurons in DR with bursting behavior. We identified 20 neurons in DR that were mostly active during wake bouts and off during NREM and REM sleep (A) and (D). These neurons, whose average waveform is demonstrated by the solid black line in (B) – standard deviation is shown in light gray shading – had bursting behavior with very short inter- spike intervals (C). The gray and orange bars (D) indicate dark and light periods respectively. We did not find any consistent or significant difference in the neurons’ firing rate during light vs dark periods.

The remaining wake-active neurons in DR were divided into three subgroups based on their average waveforms and ISIs: Most of the neurons (n=36) showed a broad asymmetric biphasic spike waveform with large positive peak and long spike duration (Fig. 2A&B). The descending portion of the waveform displayed a notch and was then followed by a smaller amplitude and longer negative component. The second type of cells (n=16) displayed a similar biphasic pattern however their spike waveforms were more symmetric and with a much less pronounced notch (Fig. 2C&D). The third type (n=10) displayed a biphasic spike waveform with a smaller positive peak which descended into a large amplitude negative trough (Fig. 2E&F). These waveforms are consistent with previous reports demonstrating wake-active DR neurons with various average waveforms both in anesthetized and un-anesthetized (head-fixed) animals (Urbain et al., 2006; Sakai, 2011).

**Figure 2.**
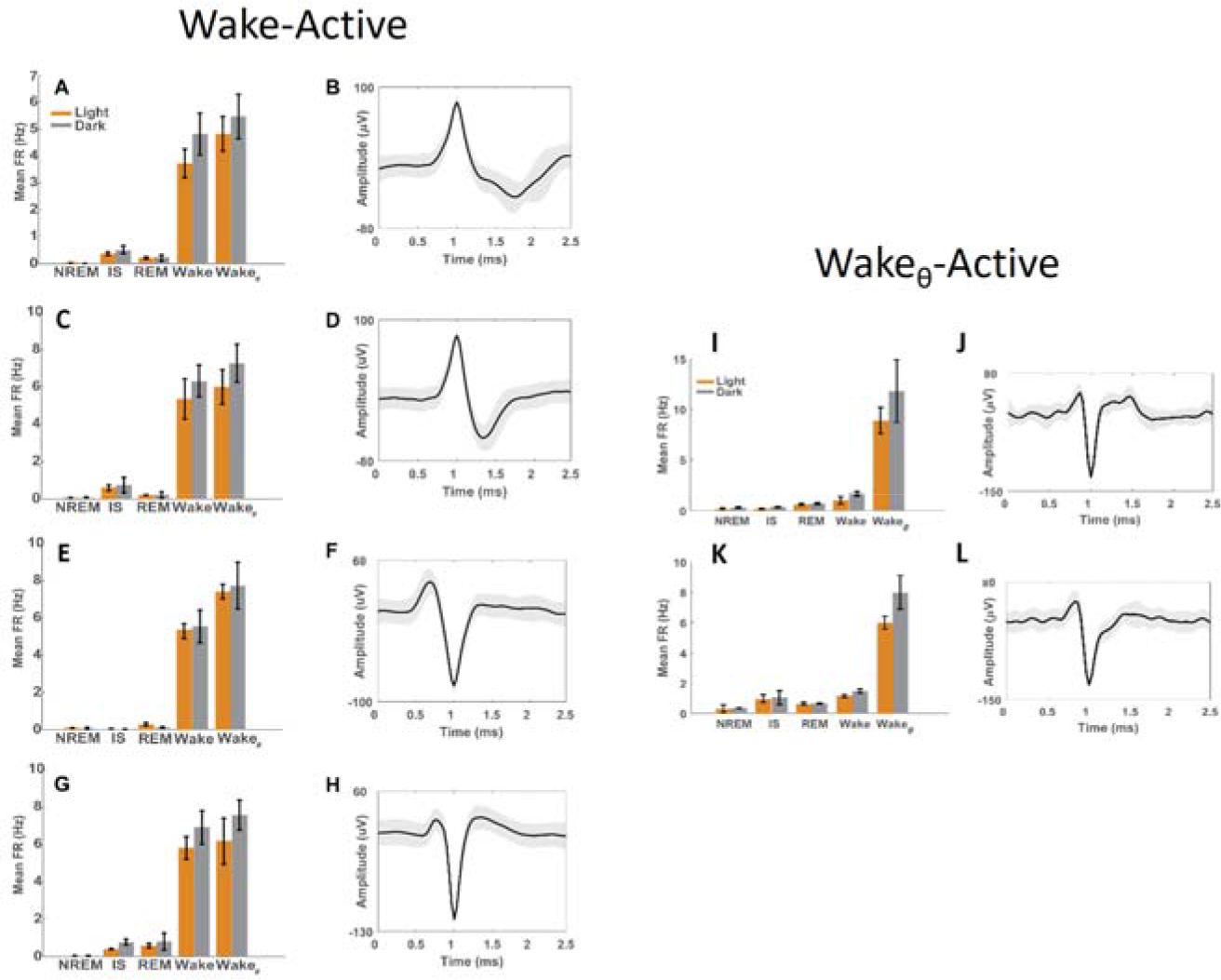
Wake and wake_θ_ active neurons. We identified 4 types of wake-active neurons in DR and 2 types of wake_θ_ active neurons in PPT and LDT. The average waveform and firing rates of the wake-active neurons (B, D, F, H) is similar to previous reports (Sakai, 2011). The gray and orange bars (A, C, E, G, I, K) indicate dark and light periods respectively. Although the neurons had higher firing rate in the dark vs light period, given the standard deviation, the differences were not statistically significant.

In 8 out of 12 rats we identified wake-active neurons in the PPT areas. These neurons displayed a triphasic spike waveform with mean spike duration of 1.36 ± 0.14 ms (Fig. 2G&H). We also identified a group of neurons in PPT (10 out of 12 rats, Fig. 2I&J) and LDT (3 out of 7 rats, Fig. 2K&L) that were only active during wake_θ_ bouts. Based on their average waveform, ISI, and state-dependency they potentially belong to the group of previously reported wake-active neurons in PPT and LDT areas (Datta and Siwek, 2002; Saper et al., 2010).

#### 3.1.2. NREM-active neurons

We identified 15 out of the 428 state-dependent neurons (1.7%), specifically in the VLPO region, with higher discharge rates during NREM than during REM or wake bouts. The activity of these neurons was continuously monitored for at least 24 hours in each animal (n = 5). These neurons had a biphasic spike waveform with a small amplitude but long negative trough (Fig. 3A&B).

**Figure 3.**
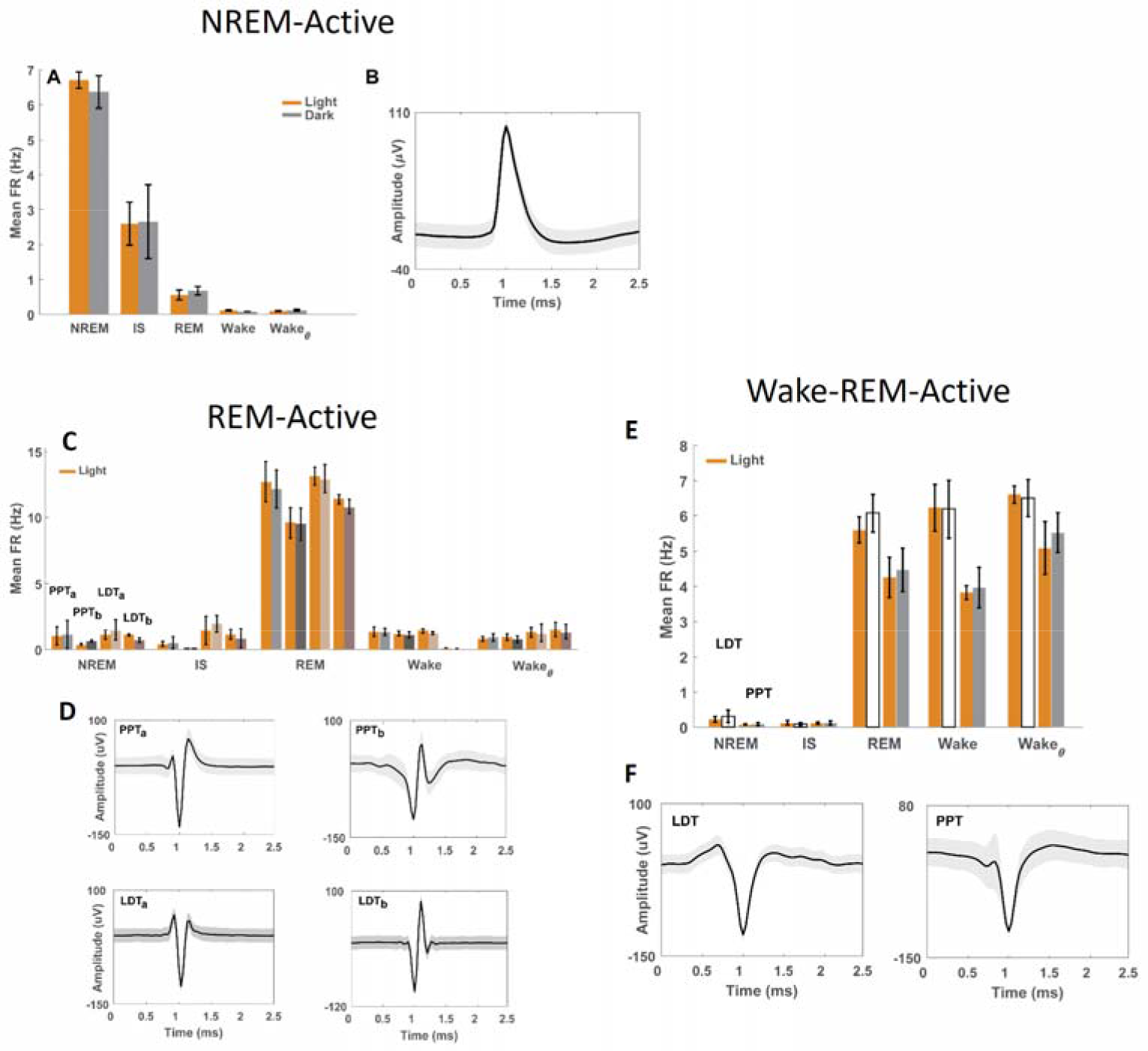
Sleep and wake-REM active neurons. We identified 15 neurons in the VLPO that were predominantly active during NREM (A, B). The average waveform (B) of these neurons is consistent with previous report (Takahashi et al., 2009; Alam et al., 2014). Similarly the activity pattern (D) and waveform (C) of the REM-active neurons in PPT (grouped as PPT_a_ and PPT_b_) and LDT (grouped as LDT_a_ and LDT_b_) were consistent with findings of Datta et al (Datta and Siwek, 2002). We also identified neurons in the LDT and PPT areas that were active during REM, wake, and wake_θ_ bouts (E). The orange bars (A, C, E) indicate light periods. The other shades of color (i.e. gray, brown, white) indicate the dark period for different neurons. We did not find any consistent or significant difference in the neuron’s firing rate during light vs dark periods.

#### 3.1.3. REM-active neurons

We identified 152 out of the 428 state-dependent neurons as REM-active. These neurons were located either in PPT (89 neurons, n=10 animals) or in LDT (63 neurons, n=7 animals) with triphasic sharp waveforms. Based on their ISI, average waveform, and location of clusters in the principle component space during spike sorting we subdivided these neurons into 2 groups for each region (Fig. 3C&D).

#### 3.1.4. Wake-REM-active neurons

A subset of recorded neurons (7%) from PPT and LDT were active both during REM and awake bouts (Fig. 3E&F). These neurons were co-localized with a portion of REM-active neurons in PPT and LDT. Therefore, differences in their activity profiles do not stem from their specific location.

#### 3.1.5. REM-off neurons

In 4 animals, we identified a number of neurons in and around the vlPAG region that were active during all SOVs except REM. The firing rate of these neurons during wake_θ_ was significantly higher than during REM (Fig. 4). Therefore, we hypothesize that although active during wake_θ_ bouts, these neurons do not code for theta rhythm or other cortical and behavioral signs of REM.

**Figure 4.**
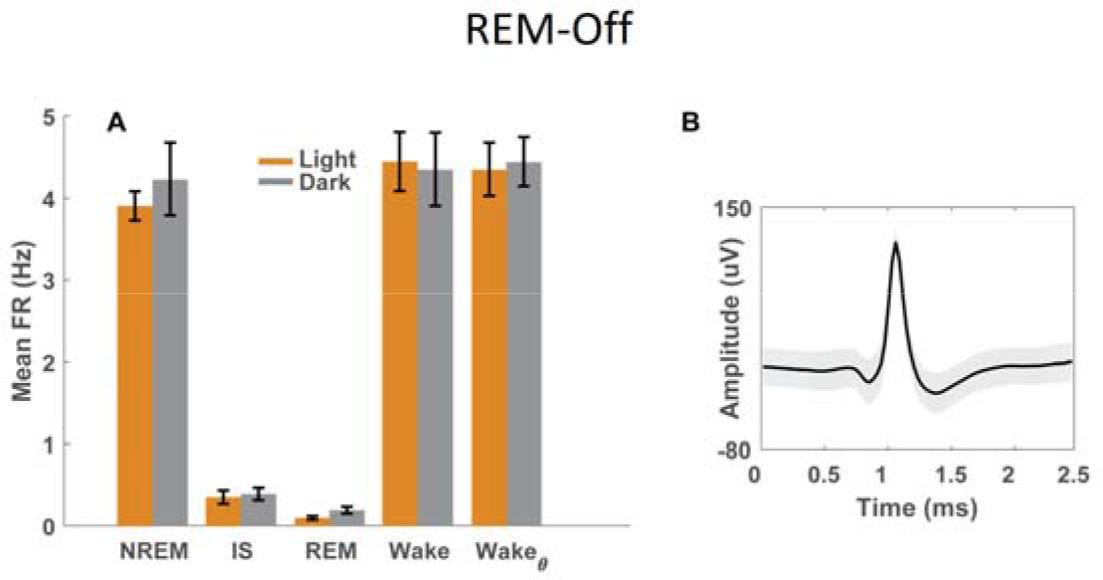
REM-off neurons. In 4 animals we found neurons potentially located in vlPAG that were active during all sleep states except IS and REM (A). These neurons did not demonstrate any theta-dependent behavior because although they are active during wake_θ_, they were significantly less active during REM (high theta activity). The gray and orange bars (A) indicate dark and light periods respectively. We did not find any consistent or significant difference in the neuron’s firing rate during light vs dark periods. The average waveform of these neurons was triphasic and similar to the cholinergic REM-active neurons in PPT and LDT.

#### 3.1.6. IS-active neurons

None of the identified state-dependent neurons active during NREM, REM, or wake states were active during the IS. However, in 4 animals we identified a group of neurons potentially within the medial preoptic area (MPA) dorsal to the VLPO that were active during REM and IS (Fig. 5). Although their firing rate was slightly lower in IS bouts that ended in REM than the ones that ended in wake, the difference was not statistically significant.

**Figure 5.**
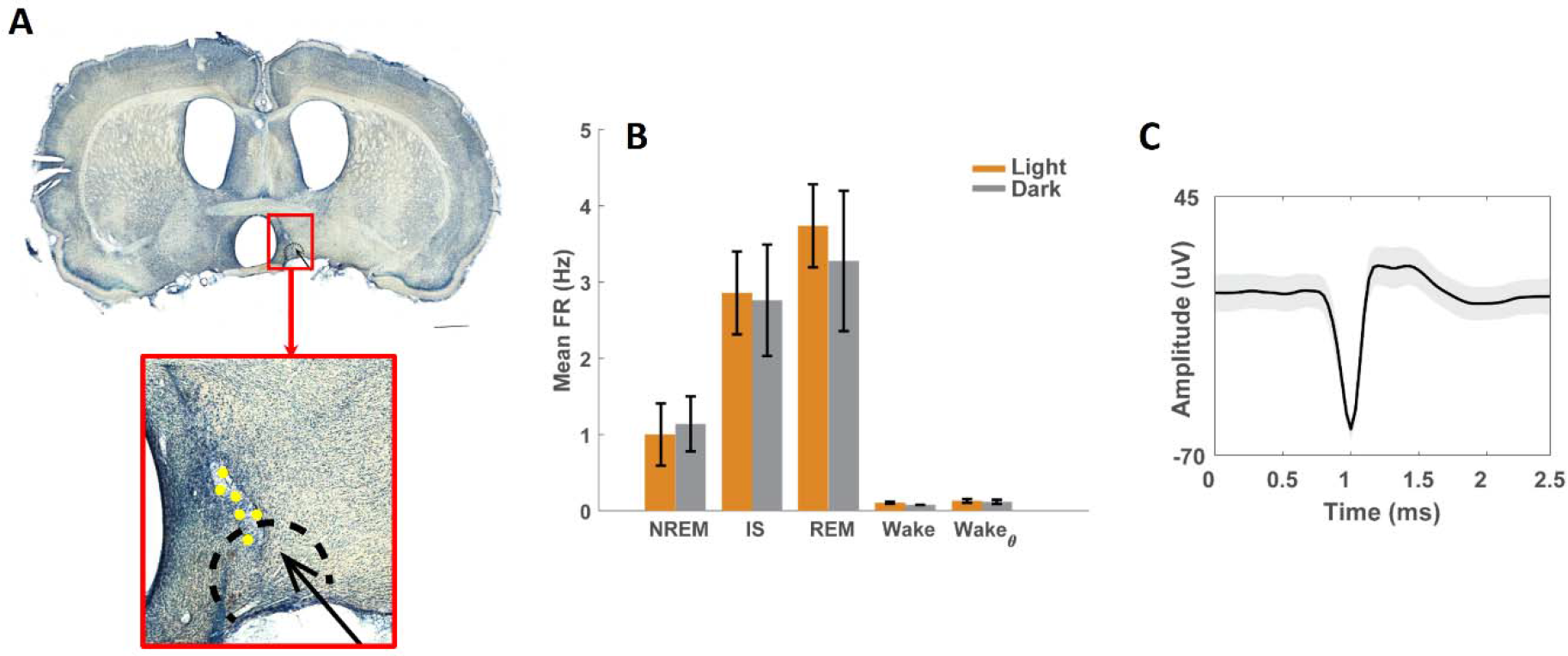
Identified neurons in the preoptic areas are active during the intermediate state. **(A)** Stereotaxic coordinates of the regions with IS-active neurons, recorded in 4 animals, are indicated by the yellow dots. For visual clarity purposes, the coronal histologic section show here is from 1 of the 4 animals chronically implanted with this target. The yellow dots indicat the electrode locations in all 4 animals when the IS-active neurons where monitored. **(B)**The IS active neurons are active during REM and IS bouts and increase their firing rate slightly upo transitions from IS to REM. However they are off during wake and wake_θ_ bouts. This indicate that 1) IS is a distinct state of vigilance with its own corresponding neuronal activity and 2) th IS-active neurons do not code for theta-rhythm activity. **(C)**Average waveform.

The IS emerges during transitions out of NREM with clear behavioral and cortical signs (see methods, Fig. S1) that, when reported (GOTTESMANN, 1996; Gervasoni, 2004; Boucetta et al., 2014), are associated with human NREM-II. Although in most reports IS appears to occur only between NREM and REM states, we found that 20% of the time IS was directly followed by wake bouts (Fig. S1C&D). Additionally, the median duration of IS in our animals was ∼45s (Fig. S2B).

The identification of neurons whose firing rate is significantly less during NREM and onl increases during transitions out of NREM into IS or REM indicates that there is a distinct circuitry that drives IS. That the IS-active neurons are active during IS and REM states, but not wake_θ_ bouts indicates that IS-active neurons are not required for coding of theta band activity.

The mean firing rate of each identified SOV-dependent neuron during each SOV is summarized in Fig. 6. The firing rates for each neuronal group (rows) during different SOVs (columns) are separated based on light (orange dots) and dark cycles (blue dots). For each neuronal group the firing rates are normalized to the maximum observed firing rate of that group across all SOVs (M_FR_, shown in black dots). The neuronal groups are separated by a combination of anatomical location and SOV-dependence, regardless of differences in their waveform patterns. For example all the identified PPT REM-active neurons (Fig. 3C&D) are grouped together. We found that although different in their waveforms, the neurons had similar activity profiles (i.e. mean firing rate, and light/dark patterns).

The data presented here pertains to the state-dependent neurons in the targeted SWRN. We aimed to investigate the original flip-flop model of sleep-wake regulation which identifies certain regions as only active during certain SOVs (i.e. NREM-active neurons in VLPO) (Saper et al., 2001; Diniz Behn and Booth, 2010). Thus, although most of the targeted regions contain multiple populations of neurons with diverse state-firing dependency, the neuronal activity presented here specifically includes NREM-active subpopulation of VLPO, REM and wake-REM active subpopulations of PPT and LDT, wake-active subpopulation of DR, REM-off subpopulation of vlPAG, and IS-active neurons that are potentially located in medial region of the preoptic area.

We found that the population mean and general activity profile of these state-dependent neurons were similar to previous reports (Datta and Siwek, 2002; Sakai, 2011, 2018; Alam et al., 2014; Boucetta et al., 2014). We observed some level of variability in mean firing rates for each neuronal group (indicated by green stars in Fig. 6) during light or dark periods. However, the variance of the firing rate distributions, specially the M_FR_ (black dots) was much lower compared to what is reported in previous studies. The statistics of these distributions are highly affected by how long and over how many SOVs each neuron’s activity is monitored. Because we have such a large sample population for each state, and for each neuron, we have a longer (and more stable) observation window over the behavior of the neuron which inherently reduces the variance. This can in turn explain the differences in distribution width compared to previous work.

Of the represented neuronal groups, VLPO is one of the most challenging structures for continuous stable recordings. This explains the dearth of reports of continuous electrode recordings, and the fact that such recordings, when available, are performed in short bouts (Szymusiak et al., 1998) or when the animal’s movements are restricted (Takahashi et al., 2009; Alam et al., 2014). To our knowledge, we are the first to report consistent multi-day continuous observations of VLPO neuronal activity. The large number of NREM bouts wherein VLPO neurons were highly active enabled calculations of firing rate with low margin of error and provided sufficient power to compare neuronal activity over different SOVs.

We found a slight difference in the mean state-dependent firing rate of a subset of neuronal groups during light versus dark periods. These included the wake-REM active neurons of PPT and LDT and the REM-off neurons of vlPAG. However, this circadian difference was neither statistically significant nor consistent in all groups. This is in line with our previous work showing no clear difference in neurons’ ISI and average waveform in dark versus light periods (Fig. 9 in (Billard et al., 2018)).

### 3.2. Investigation of causality of neuronal activity in emergence of cortical signs of SOV

Our aim was to investigate the role of state-dependent neurons, within the conventional brainstem-hypothalamic flip flop model of sleep-wake regulation, in the emergence of SOVs and SOV transitions. There is a tendency to implicitly interpret the term ‘XXX-active neuron’ (i.e. REM-active) as a neuron whose activity is causal of initiating the ‘XXX’ state (i.e. REM state). An alternative role denoted in the literature is that such neurons are involved in the maintenance of the state in which they are active (i.e. (Grace et al., 2014; Xu et al., 2015; Weber and Dan, 2016; Sakai, 2018)). We therefore investigated whether changes in firing rate could be causal in state initiation – as indicated by significantly increasing firing rate ahead of the observed transition time.

Neuronal firing rates over many sleep cycles (∼100-300) were derived from simultaneous measurements of single-unit activity from a subset of the SWRN. For each SOV transition (from state A to B), we collected the distribution of firing rates while in state A. The firing rate was calculated from spike-times away from state transitions by at least 3 seconds. Procedurally, we determined SOV transition times (t_trans_) as the time when EEG and behavioral signs of state B first emerged. Such signs emerge from brain regions downstream from the SWRN, and therefore t_trans_ as defined must be delayed from transitions in activity state of the SWRN. Changes in firing rate of a particular neuron or neuron type that follow t_trans_ cannot be causal for the transition, while those that precede it can but are not guaranteed to be so.

Following the Poisson process statistics (see Methods) we determined the time-point that the instantaneous state-conditioned firing rate during state B differed significantly from the firing rate in state A. The time-point is denoted as t_sig_. The large ensemble of firing rates for each neuron during each transition allows for sub-second resolution determination of t_sig_.

If the significant change in the state-conditioned firing rate occurred prior to transition time t_sig_ < t_trans_, then it is counted as negative t_sig_. Positive t_sig_ indicates that the state-conditioned firing rate changed after transition time, t_sig_ > t_trans_. Therefore, a positive t_sig_ implies that the changes in firing of that neuron type **cannot** be causal for the state transition. In addition, our sleep scoring was done with 1s discretization. Therefore, there may be as much as an average of 0.5s delay in marking t_trans_ from the actual change in signs. This means that an even more conservative scoring to which changes in neuronal firing could be causal would be t_sig_ < t_trans_-0.5.

The t_sig_ values for the 428 identified state-dependent neurons are marked by blue dots in Fig. 7. The gray shading indicates the positive t_sig_ and therefore non-causal relationships. The white shading indicates the negative t_sig_ where the change in firing rate occurred prior to state transition. For each cell group and state transition, dashed gray lines indicate the representative average state-dependent firing rate as a function of time during state transitions. We found that the t_sig_ values for individual cells are tightly distributed within each cell group for each state transition. This indicates the consistency of neuronal activity during state transitions.

Cell groups whose mean firing rates increase during state transitions (e.g. REM-active PPT or LDT neurons during NREM to REM transitions) can either turn on to cause the state (t_sig_ < 0, white shading in Fig. 7), which means high discharge rates prior to emergence of cortical signs of a SOV, or can be turned on as a consequence of state transition (t_sig_ > 0, gray shading in Fig. 7). Among all cell groups with increasing firing rate during state transitions, only NREM-active neurons of VLPO and REM-off neurons of vlPAG during REM to NREM transitions had negative t_sig_. The discharge rate of all other neurons increases *after* the appearance of EEG and behavioral signs of SOV. Therefore, the increase in their discharge rate does not mean that they are causing the state change, only that their behavior is a consequence of the state change.

Likewise, cell groups whose mean firing rate decreases during state transitions (e.g. wake- active DR neurons during wake to NREM transitions) can either be responsible for the emergence of the state (t_sig_ < 0, white shading in Fig. 7), or be inhibited as a consequence of the state (t_sig_ > 0, gray shading in Fig. 7). We found that all the cell groups that decreased their firing rates during state transitions did so prior to transition time (t_sig_ < 0), which could allow a new SOV to emerge. We anticipate that such inactivation is sourced from changes in the activity of the presynaptic source.

The NREM-active neurons of VLPO are the only population with negative t_sig_ values during all state transitions. Specifically, during transitions into NREM the activity of the VLPO neurons increases significantly with respect to the state-conditioned average firing rate prior to the state transition. These neurons also decrease their firing rate significantly prior to transitions out of NREM. Therefore, the causality hypothesis (Szymusiak et al., 1998; Saper et al., 2005, 2010) holds for the NREM-active neurons of VLPO.

### 3.3. Neuronal activity in preoptic area and vlPAG demonstrate distinct dynamics during IS

Because we found that IS seems to be characteristic of ending of NREM and often precedes REM, we utilized the t_sig_ values to investigate the behavior of REM and NREM active neurons during IS transitions. During NREM to IS transitions, the discharge rates of both the solely NREM-active neurons of VLPO and the REM-off neurons of vlPAG decreased prior to transition into IS (t_sig_ < 0). The behavior of NREM-active neurons implies a level of causality, but whether the decrease in firing rate is programmed internally or caused by other factors remains to be investigated. In contrast, the discharge rate of the REM-active as well as REM-wake-active neurons of PPT and LDT does not change during NREM to IS transitions.

The IS-active neurons remained active with no substantial change in their firing rate during IS to REM transitions. However, we observed a statistically significant decrease in their firing rate during IS to wake transitions (t_sig_ < 0). Similarly these neurons stopped firing prior to transitions out of REM to NREM or wake (t_sig_ < 0). These indicate that IS-active neurons might be coding for transitions out of IS (or REM).

The activity patterns of the NREM-active neurons in VLPO, REM-off neurons in vlPAG, and IS- active neurons in the preoptic area provide another line of evidence that IS is a distinct state of vigilance rather than only a transitory dynamic between NREM and REM.

## 4. Discussion

We developed a dataset including 569 cumulative days of SOV-scored single- and multi-unit measurements from multiple cell-groups in the sleep-wake regulatory network (SWRN). These measurements are from 12 freely behaving rodents, with each rat recorded for over 30 day periods, which consisted of multiple 3-10 day long segments of continuous recordings. The dataset (previously presented in (Billard et al., 2018)) includes unit activity from up to five noncoplanar, noncollinear targets of the SWRN (PPT, LDT, DR, vlPAG, VLPO, and MPA), along with ECoG and LFP recordings of cortex and hippocampus in freely behaving rats.

Current theory about wake-sleep regulation posits that there are wake-promoting and NREM-promoting cell groups, and that these inhibit one another. This theory would predict changes in the wake-promoting groups before awakening from NREM sleep, and in the NREM-promoting neurons prior to transitions into NREM sleep. Our dataset complements previous work testing this theory (Aston-Jones et al., 2001; Datta and MacLean, 2007; Saper et al., 2010; Takahashi et al., 2010; Sakai, 2011; Alam et al., 2014; Boucetta et al., 2014; Van Dort et al., 2015; Cox et al., 2016; Weber and Dan, 2016), but is characterized by several significant distinctions.

First, our simultaneous measurements of hippocampal LFP, ECoG, head acceleration, and neuronal activity from multiple cell-groups in freely behaving animals over long (3-10 day) continuous recording periods in the animals’ home cages represent a novel “systems neuroscience” attempt aimed at characterizing the dynamics of the brainstem-hypothalamic SWRN system described by Saper et al. (Saper et al., 2001, 2010). Second, the data include continuous and simultaneous measurements from multiple cell groups of SWRN (including

NREM-active VLPO neurons) over many (100-300) consecutive sleep-cycles without introduction of any restrictions to affect animal behavior. Third, our approach to sleep scoring resulted in continuous and temporally precise determination of SOV and SOV transition times. Fourth, time-resolved analysis of the SOV and SOV transition times as well as acquisition of sufficient sleep-wake cycles provide suitable statistical power to probe causality between neuronal firing rates and emergence of cortical signs of states. Finally, we identified a subset of neurons, potentially located in or close to the medial preoptic area (MPA) that code for the intermediate state (IS, (GOTTESMANN, 1996; Boucetta et al., 2014)). We therefore characterized IS as a distinct sleep state from NREM that dominantly transitions to REM.

Much of the research investigating the neuronal activity of the cell-groups in the SWRN is performed within conditions that inherently modulate the animals’ sleep cycles and/or limit the time-resolution required for determination of the relationship between firing rate and state transitions. Almost all sleep studies available today are performed with head-fixed animals (Takahashi et al., 2006, 2010; Sakai, 2012, 2018), or in novel environments such as isolation boxes, and provide short recordings of the sleep cycles (Szymusiak et al., 1998; Datta and Siwek, 2002; Takahashi et al., 2006; Sakai, 2011, 2012). While such methods avoid the technical challenges associated with simultaneously recording the hippocampal, cortical, and behavioral indices of different sleep states as wells as state-dependent neuronal activity, they complicate the analysis of SWRN dynamics.

Deviations from normal home-cage environments change the animals’ normal behavior and sleep-wake cycles. Short-term recordings from restricted animals eliminate the effect of exogenous (light-dark cycles) and endogenous circadian cycles on sleep dynamics. Furthermore, to detect a statistically significant change in neuronal firing rates during state transitions, with sub-second temporal resolution, many sleep cycles are required. Finally, conventional SOV scoring techniques assign a score to non-overlapping time-windows (Takahashi et al., 2009, 2010; Bastianini et al., 2017; Sakai, 2018; Grieger et al., 2021; Huffman et al., 2021) with fixed length and predetermined boundaries. These fixed windows prevent accurate assessment of the SOV dynamics and identification of the true onset of a state.

### 4.1. Implications of the intermediate state for sleep-wake regulation

Gottesmann was the first to identify an intermediate state in Wistar rats that occurred between NREM and REM, and which exhibited both cortical spindles (a sign of NREM) and hippocampal theta rhythm (a sign of REM) (Gottesmann, 1964) . Gottesmann’s finding of IS was further confirmed both in later rat experiments (Gandolfo et al., 1990; Neckelmann et al., 1994; GOTTESMANN, 1996) and observations of human sleep (Lairy, 1966). More recently this state has appeared in a small subset of publications focused on sleep-wake regulation in healthy (Gervasoni, 2004; Boucetta et al., 2014) and diseased brain (Andrade et al., 2017; Pitkänen et al., 2019). We also identified IS in our animals and found distinct temporal, spectral, and neuronal patterns associated with it.

The conventional/routine sleep-scoring analyses assume state transitions are discrete and easily classified by fast changes in EEG features, and in behavioral and physiological activity represented by EMG power, head acceleration, heart rate, and/or breathing. Therefore, research has focused on identifying brain regions involved in triggering discrete transitions between these states. However, the existence of a state with clear spectral and neuronal characteristics that combines and expands upon the features of both NREM and REM in the transition from NREM to REM or to wakefulness poses new questions regarding the mechanisms underlying the onset of any SOV.

The IS-active neurons reported here remained active during IS to REM transitions. However, these neurons turned off prior to transitions out of IS and into wake. Therefore, we hypothesize that the activity of these neurons determines whether transitions out of IS progress into REM or wake. Based on our histological analysis, these neurons might be located in the medial regions of the preoptic area.

The technical challenges in measuring from a number of SWRN indicate the value of SWRN mathematical models (see for example (Tamakawa et al., 2006; Fleshner et al., 2011)) for understanding the underlying mechanisms of sleep-wake regulation. However, the models and the experiments they are based on have generally not considered the existence of the IS or the unique sets of IS-active neurons. Implementation of the IS as a distinct SOV with its own state-active neurons will enable the models to provide mechanistic insight into differentiation between transitions to wake versus transitions to REM.

### 4.2. Implications of correlative vs. causal activity in the brainstem-hypothalamic SWRN

We used time-resolved transition-based analysis to probe the causality of the SWRN dynamics in transitions between states of vigilance. With the exception of NREM-active VLPO neurons, increases in state-conditioned mean firing rate always followed the emergence of the state which is determined from cortical and behavioral signs. A change in neuronal activity that follows the SOV cannot be programming it.

Previous work including brief recordings (Datta and Siwek, 2002) and lesion experiments (Saper et al., 2001) has indicated that PPT neurons are involved in both the induction and the maintenance of REM from NREM sleep and are a precursor to the signs of REM (Datta and MacLean, 2007). We too observed increases in PPT activity during REM. However, we observed increased PPT activity only *after* emergence of cortical signs of REM. This questions the hypothesized role of cholinergic REM-on PPT neurons in REM induction. We suspect that this mismatch is due to two factors. First, the spectral and temporal features used in our sleep-scoring are generated via causal filters and updated every second at the end of computation window. Therefore, all the scores are based on information up to the time leading to the marked state transition and *not* after that. This is in contrast to fixed length windows used in conventional sleep-scoring that are susceptible to missing the correct transition time, or hand-scored versions that use a-causal potentially biased methods to estimate transition times. Second, for each neuron we collected a large ensemble of firing rates computed over many sleep-wake cycles. The large ensemble size enables sub-second resolution in determining peri-transition-time changes in firing rate (see Methods).

We observed a similar behavior for other identified SWRN nuclei. Neurons in REM-active LDT, Wake/REM-active PPT and LDT, Wake-active DR, Wake-active PPT, and REM-off vlPAG cell groups all show a significant change in their firing rate only *after* their corresponding state transitions.

The cell groups involved in sleep-wake regulation are functionally and spatially heterogeneous and extend across large neural territories. Many aspects of how such a large number of variable cell groups interact to govern rapid ultradian transitions within a sleep cycle (i.e. NREM to REM) as well as much slower homeostatic and circadian processes are still unknown. Acute disruptions of the interactions within the SWRN via anatagonist injections or optogenetic inhibition are shown to cause acute loss of the corresponding SOV. However, chronic ablation of the basal forebrain cholinergic neurons (Kaur et al., 2008), the tuberomammillary histaminergic neurons (Gerashchenko et al., 2004), the LC and pontine cholinergic neurons (Webster and Jones, 1988; Shouse and Siegel, 1992; Lu, 2006), or combinations of these structures (Blanco-Centurion et al., 2007) have minimal effects on the regulation of that SOV.

Further, almost all of the SWRN contain intermingled cell types that are active during more than one SOV. For example in the basal forebrain, glutamatergic, cholinergic, and parvalbumin (PV)- expressing GABAergic neurons are shown to be active during both REM and wake, but their optogenetic activation promotes only wakefulness (Xu et al., 2015). A routine based on acute study of stimulus (lesion/optogenetics)-response (i.e. changes to SOV length and occurrence) dynamics might not be fitting for investigation of a naturally occurring spontaneous behavior such as sleep-wake transition and cycling. Therefore, although internal/external stimuli might cause transitions into a state, it does not necessarily mean that the stimulated neurons are causal for the emergence of the state in normal conditions.

The *in vivo* experimental approach described here and in (Billard et al., 2018) allows for simultaneous recording of neuronal activity in multiple deep brain structures throughout many days in freely behaving animals. As evidenced by the findings reported here, this approach has the potential to uncover new insights on mechanisms underlying the dynamics of sleep-wake regulation.

## 6. Appendix

**Table S1.** Detailed recording information from brainstem-hypothalamic structures. The cell include information about which targets were recorded from successfully as well as the numbe of well-defined SOV-dependent and total units counted over the entire course of recordings fo each axis and animal. The final row is the sum of SOV and total units over all animals in the cohort. Animals 1-5 had three Microdrive axes, animals 6-11 had a total of four microdrive axes and animal 12 had a total of five microdrive axes. Formatting of cells refer to the following conditions: recorded units with histological validation that the electrodes hit the target (green) recorded units with no histological validation (orange). For this group the state-dependency of the unit-recordings was established based on EEG and behavioral measures; recorded activity but no units crossed the 7 SD threshold (red). Some implanted microwire bundles were kinke (K) during the Microdrive implant procedure or later driving sessions, which prevente electrodes from driving. Sessions were continuous recording periods between electrode-drivin sessions. Recording sessions typically lasted between 3 and 10 days.

| Animal #<br>(Sex) | Days Recorded<br>[# of Sessions] | PPT |  | LDT |  | DR | LC | VLPO |
| --- | --- | --- | --- | --- | --- | --- | --- | --- |
|  |  | Left | Right | Left | Right |  |  |  |
| 1 (M) | 41 [13] |  |  |  |  | 47 (23) |  |  |
| 2 (M) | 37 [12] | 42 (21) | 33 (13) |  |  | 43 (17) |  |  |
| 3 (M) | 57 [9] | 29 (19) | 29(8) |  |  | 18 (10) |  |  |
| 4 (M) | 73 [12] | 7 (1) |  |  |  | 18 (7) |  |  |
| 5 (M) | 66 [10] | 20 (9) |  |  | 14 (4) | 27 (10) |  |  |
| 6 (M) | 43 [8] | 24 (20) |  |  | 29 (22) | 32 (11) |  | 26 (10) |
| 7 (F) | 39 [7] |  | 6 (4) | 19 (12) |  | 3 (2) |  | 9 (4) |
| 8 (F) | 54 [9] | K |  |  | 47 (28) | 44 (31) |  |  |
| 9 (M) | 57 [9] | 8 (6) |  |  | 43 (21) | 23 (3) |  | 13 (5) |
| 10 (M) | 34 [8] |  | 42 (27) | K |  | 15 (9) |  | 16 (6) |
| 11 (M) | 29 [8] |  | 25 (17) | 30 (19) |  |  |  | 16 (5) |
| 12 (M) | 39 [6] |  | 30 (14) | 16 (7) |  | K | 4 (0) | 8 (3) |
| Total Neuron Count<br>SOV-Dependent Neurons () |  | 295 (159) |  | 198 (113) |  | 270 (123) | 4 (0) | 88 (33) |

### 6.1. Characterization of the intermediate state

**Figure S1.**
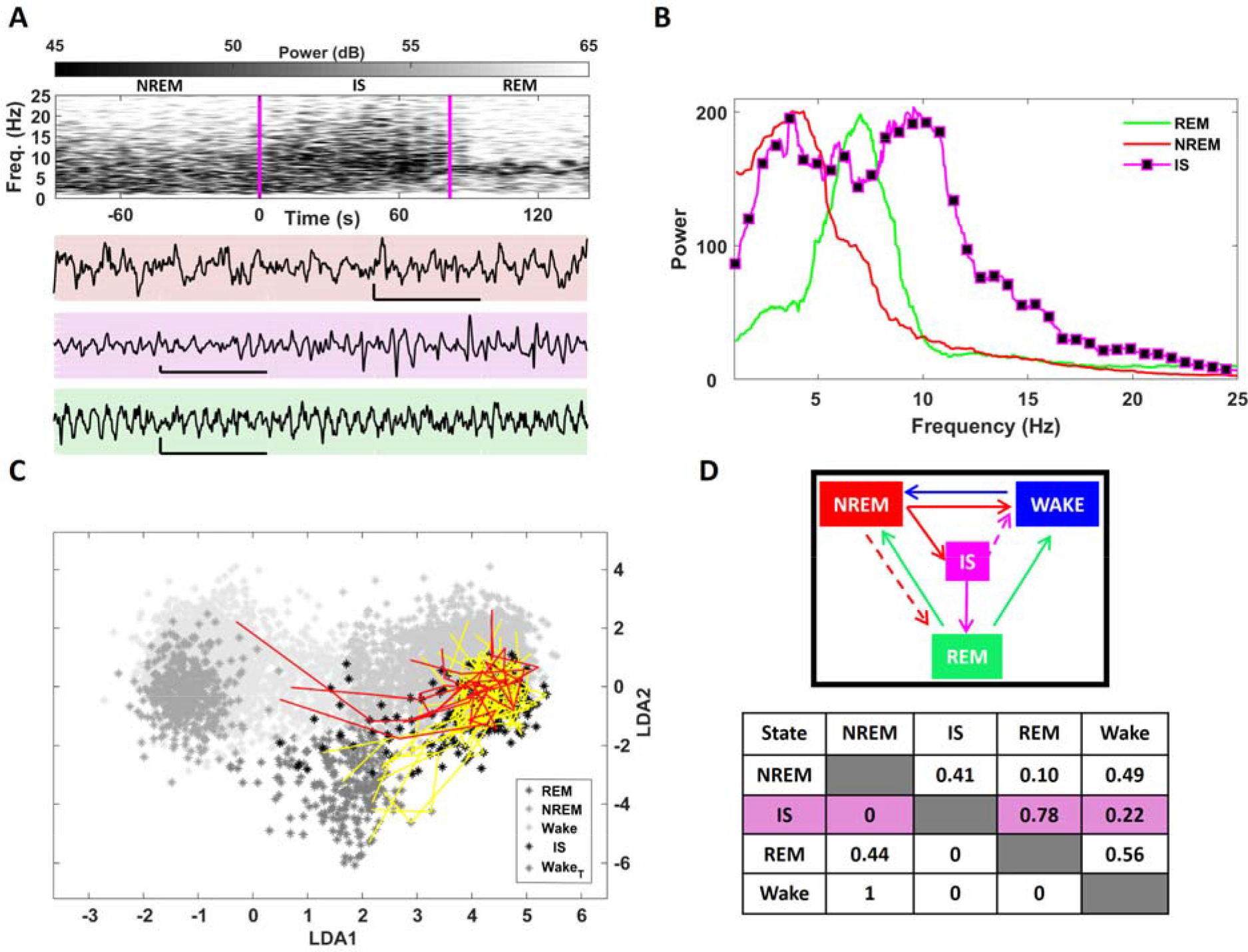
Time-frequency characteristics of the intermediate state. The IS was observed in almost all NREM to REM transitions. **(A)** In an example hippocampal LFP recording in one animal, the power spectrum during NREM shows lower frequency activity in the delta range (high delta peak in red in B). The transition to REM is not discrete. As the brain exits NREM, power increases in the theta range appear simultaneous with the persisting peaks in the delta band (theta and delta peaks in magenta in B). The IS onset and end are marked by the solid magenta lines. The REM onset is then marked by a clear consistent peak in the theta band (high theta peak in green in B). Example time-series traces for NREM (red panel, ECoG), IS (magenta panel, LFP), and REM (green panel, LFP) are shown in the bottom panels. All vertical and horizontal scale bars are 100 mV and 1 second; respectively. Power is calculated over 8 second windows with 4 second overlap. **(B)** The simultaneous delta and theta activity in the IS is highlighted in the power plot averaged over 20 second long time-series for each state. Power is calculated from the same hippocampal LFP recording as in A over 2 second long windows with 1 second overlap. **(C)** Scatter plot of the first two linear canonical discriminants highlight the three main groups. Each dot corresponds to a 2 second window for which the spectral features were calculated. For clarity, the dataset presented here is from 48 hour of recording in one normal animal. After the clustering analysis, within each cluster, the 95% most-inclusive contour was chosen as the boundary of that cluster. We define the transitions between states as trajectories connecting the clusters in the canonical space. Almost all transitions from NREM to REM (yellow) or to Wake (red) seem to first dwell in the IS region (black dots). **(D)** Shown in the upper panel are the possible transitions between SOVs. The majority of transitions from NREM to REM passed the IS region, therefore direct transitions (much fewer) were marked by a red dashed line. The transition probabilities between SOVs, across all animals, are shown in the bottom panel. The majority of transitions out of IS state end in REM (78%), whereas only 22% of those transitions end in Wake.

**Figure S2.**
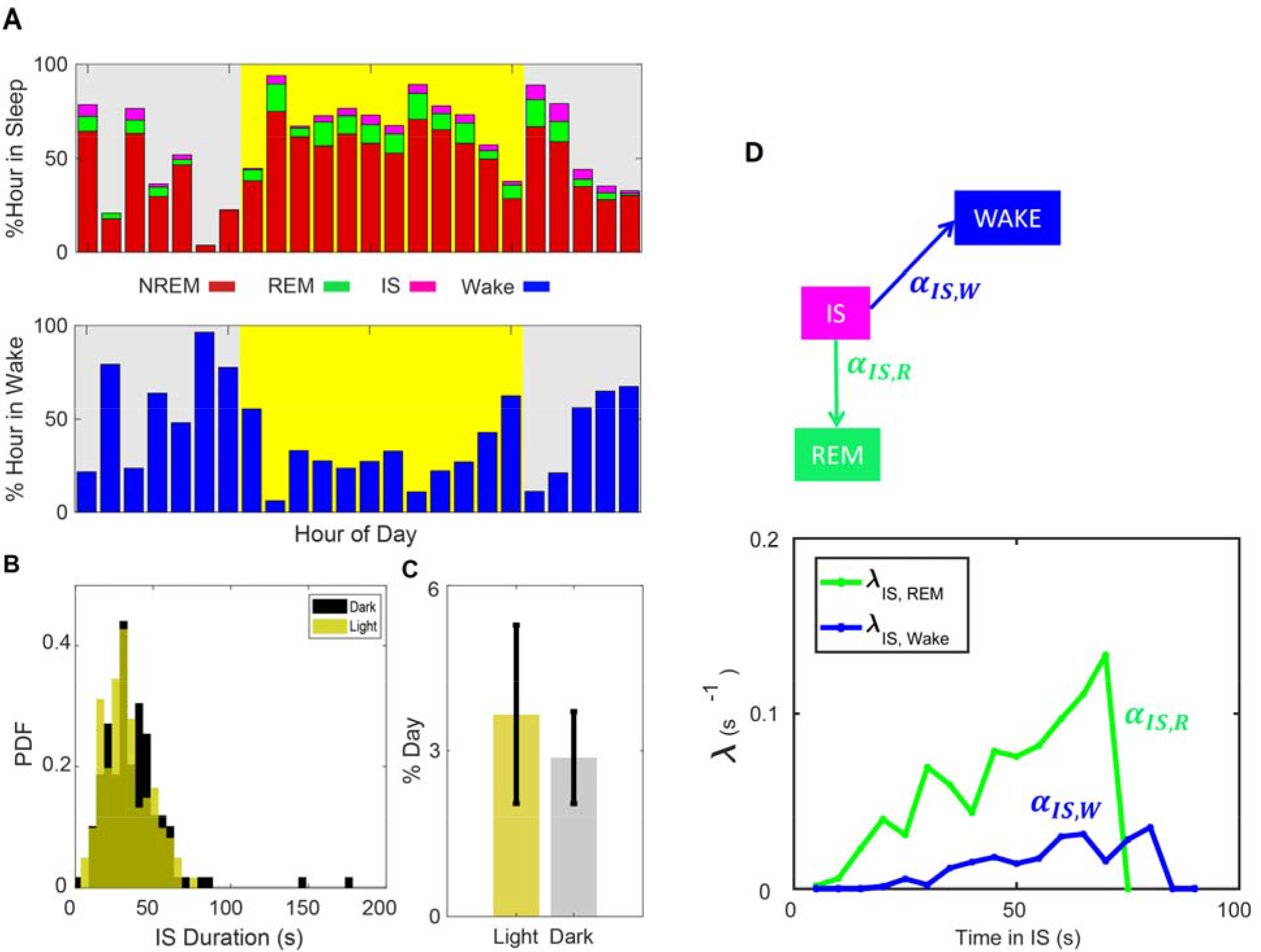
IS patterns and duration across the light/dark cycle for all rats. **(A)** Hourly percentages spent in NREM (red), IS (magenta), REM (green), and Wake (blue) are shown for all rats. The majority of the light period (yellow) is spent in sleep and the majority of the dark period (gray) is spent in Wake. **(B)** Distribution of the duration of the IS pooled from all animals both during light (blue) and dark (orange) periods. Regardless of the time of day, the intermediate state lasts for a considerable amount of time that should be detectable in the current sleep-scoring algorithms. **(C)** Mean and standard deviation of percent of day spent in IS for light and dark period. Difference in the light/dark cycle was calculated using a one-sided unequal variance t-test with alpha=0.05. Although animals exhibit an increase in daily IS time during the light cycle, it is not significant. **(D)** IS bouts pooled across all rats (n=13) were scored as either IS → REM or IS → Wake. Hazard rates were calculated using competing groups. Competing groups analysis assumes one event precludes the other. λ_REM_ is on average 4 times larger than λ_Wake_. This ratio indicates the relative frequency of REM and Wake transition types in the data set. The λ_REM_ first grows and then quickly decays, with peaks at IS duration of 30 and 70 seconds. λ_Wake_ remains comparatively flat with a small peak at 80.

### 6.2. Characterization of neural activity during state transitions

Our main goal is to understand the role of brainstem and hypothalamic neurons in the emergence of sleep-wake states and the transitions between them. By clarifying the temporal relationship between spontaneous neuronal activity and state transitions in freely behaving animals, we can elucidate the role of specific cell-groups in the initiation or maintenance of a SOV. Therefore, we acquired simultaneous measurements of single- and multi-unit activity from a subset of the sleep-wake regulatory cell-groups as well as hippocampal LFP and ECoG.

As described in the methods, SOV and state transition times were defined from the hippocampal, cortical, and behavioral signs of the state, using spectral features of the hippocampal LFP, ECoG, and head acceleration.

We first examined the activity of each identified neuron during all states of vigilance. We extracted the firing rates for the identified neurons and validated their location via histological analysis. An example of these processing and analysis steps for an identified REM-active neuron in the PPT cell-group is shown in Fig. S3. The data shown are from one animal, over all IS to REM instances (n = 168) in one 5-day recording session. The histological analysis (Fig. S3A) confirmed that the electrode was in PPT. Likewise, the neuron’s waveform (Fig. S3B) and inter-spike interval histogram (ISIH) is consistent with cholinergic cells of the PPT (Fig. S3C) (Satoh and Fibiger, 1986; Datta and Siwek, 2002; Grace et al., 2014; Van Dort et al., 2015).

Every detected spike of this particular neuron during every IS to REM transition was marked (raster plot in Fig. S3D upper panel) and the average state-conditioned firing rate, as a function of time with respect to transition time, was computed. The average state conditioned firing rate (over all REM to IS incidents) for this particular PPT neuron during both the IS and REM is shown in black traces in the bottom panel of Fig. S3D. Consistent with a REM-active neuron (one whose activity is high during REM) the average firing rate is nearly constant for times positive with respect to the transition time, which is consistent with the raster-plot having consistent firing in the green-shaded regions (Fig. S3D upper panel). Note that the variance in firing rate goes up for large time periods because there are fewer very long REM periods over which it is calculated.

To calculate changes in firing rate during state transitions, we used Poisson statistics. The pre-transition average firing rate over many trials formed the reference distribution. An example of the mean of the average firing rate distribution during the all the ISs is indicated by the solid dashed line in the lower panel of Fig. S3D. We then used the cumulative distribution function with 1%-99% confidence intervals to find the post-transition instantaneous firing rate outside of the confidence bounds. The time point associated with the significantly different instantaneous firing rate was then determined as t_sig_. The orange dashed line in the lower panel of Fig. S3D indicated the t_sig_ for this specific neuron.

In addition to the state-conditioned firing rates, we simultaneously compute and report the probability of being in a state, P(state). The green and magenta lines in the bottom panel of Fig. S1D indicate the P(IS) and P(REM). By construction, according to the homeostatic sleep drive, once the dynamics transition into a state, the probability of staying in that state monotonically decreases over time. The P(state) shown in Fig. S3D is an example of this phenomenon, where the P(REM) monotonically decreases during time spent in REM.

The procedural steps detailed in Fig. S3 for one PPT neuron were implemented to collect the state-conditioned firing rates and P(state) for all identified neurons as well as for all periods before and after every allowed transition type.

**Figure S3.**
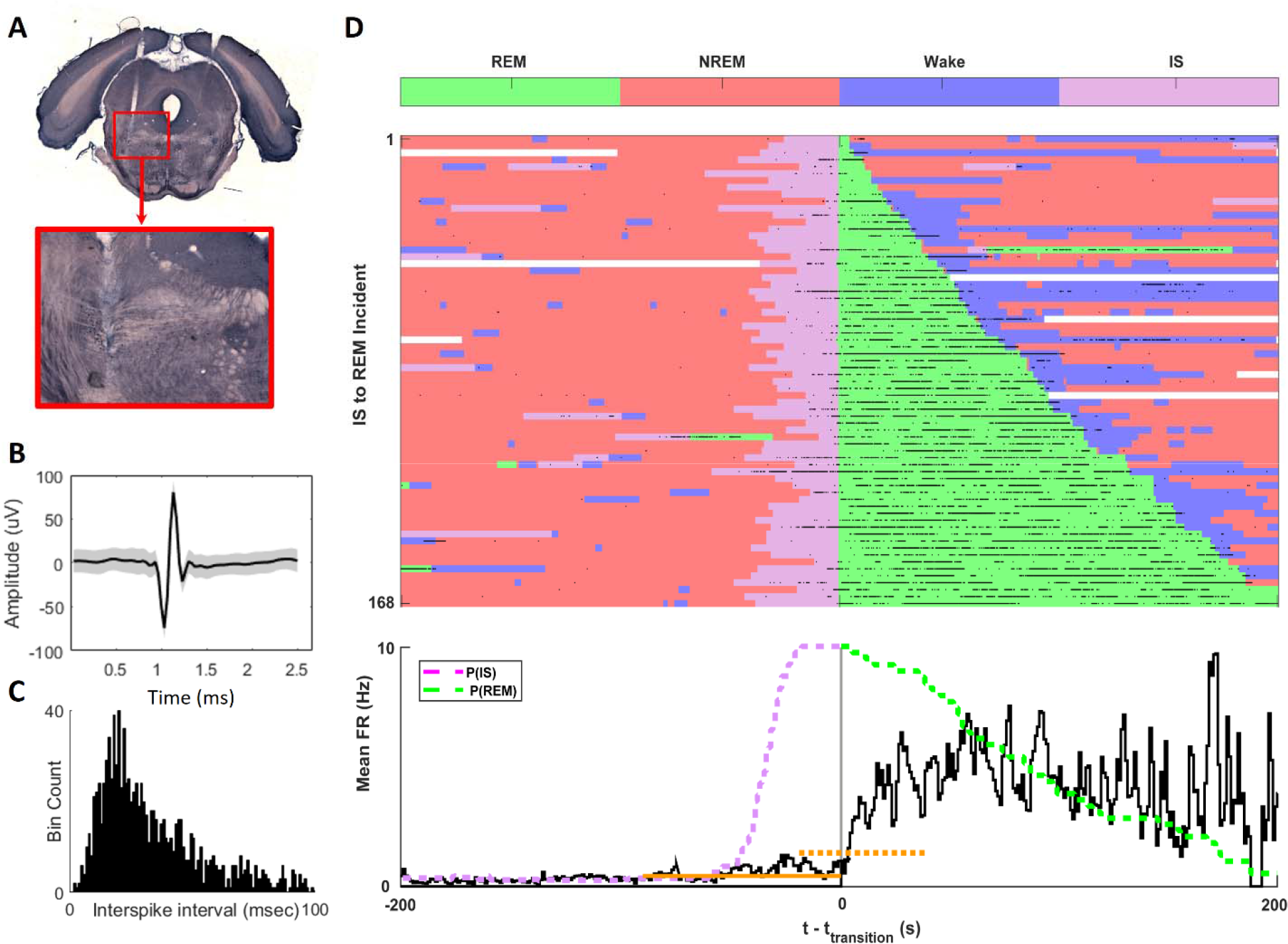
Activity analysis of a representative REM-active PPT neuron. **(A)** NADPH-stained coronal section of the brain show the cholinergic population in the PPT as well as the electrode track in the PPT. **(B)** Average waveform of the identified action potential for one PPT neuron over one 5-day long driving session. **(C)** The inter-spike interval (ISI) of the neuron. **(D)** Peri-REM firing rate of a PPT neuron in one animal. The raster plot is ranked according to the duration of REM bouts and with respect to time since transition into REM. The firing activity of the neuron is indicated by the black markers. In a raster plot format they each indicate every time an action potential is recorded. We calculated the probability of each state given time from state onset; indicated by the dashed lines in the bottom panel (magenta for IS; P(IS) and green for REM; P(REM)). On average, the PPT neuron fires much more infrequently during the IS bout, and only significantly increases its activity during REM (green area). The significance is calculated based on the Poisson statistics, where the reference firing rate distribution is the average firing rate of the prior state (here, IS) over many trials indicated by the solid orange line. The time point when the instantaneous firing rate falls out of the 1%-99% confidence bounds is marked as t_sig_ marked here by the orange dashed line. As P(REM) decays, so does the average firing rate. The average firing rate here indicates the neuronal activity conditioned on state and is calculated in 1 second bins. The probability traces are rescaled from [0,1] to [0,10] to match the scale of the average firing rate.

